# Locality-sensitive hashing enables signal classification in high-throughput mass spectrometry raw data at scale

**DOI:** 10.1101/2021.07.01.450702

**Authors:** Konstantin Bob, David Teschner, Thomas Kemmer, David Gomez-Zepeda, Stefan Tenzer, Bertil Schmidt, Andreas Hildebrandt

**Affiliations:** Institute of Computer Science, Johannes Gutenberg University, Mainz; Institute for Immunology, University Medical Center of the Johannes Gutenberg University, Mainz

## Abstract

Mass spectrometry is an important experimental technique in the field of proteomics. However, analysis of certain mass spectrometry data faces a combination of two challenges: First, even a single experiment produces a large amount of multi-dimensional raw data and, second, signals of interest are not single peaks but patterns of peaks that span along the different dimensions. The rapidly growing amount of mass spectrometry data increases the demand for scalable solutions. Existing approaches for signal detection are usually not well suited for processing large amounts of data in parallel or rely on strong assumptions concerning the signals properties. In this study, it is shown that locality-sensitive hashing enables signal classification in mass spectrometry raw data at scale. Through appropriate choice of algorithm parameters it is possible to balance false-positive and false-negative rates. On synthetic data, a superior performance compared to an intensity thresholding approach was achieved. The implementation scaled out up to 88 threads on real data. Locality-sensitive hashing is a desirable approach for signal classification in mass spectrometry raw data. Generated data and code are available at https://github.com/hildebrandtlab/mzBucket. Raw data is available at https://zenodo.org/record/5036526.

## Background

### Mass spectrometry in proteomics

Valuable information for medicine and design of new drugs for several severe diseases [1] are expected to be gained by new discoveries in proteomics [2][3][4], the field that studies proteins experimentally on a large scale. An experimental technique commonly used in proteomics is mass spectrometry (MS) [5], which allows to separate ionized molecules by their mass-to-charge ratio (m/z) and which can be combined with measurement of other physical and chemical properties.

The overall goal in an untargeted MS-based proteomics experiment is to identify and quantify as many proteins as possible in a given sample with a high quantitative performance in terms of precision and reproducibility. In particular, in bottom-up proteomics proteins are digested into peptides first and then those peptides are measured.

If only the mass-to-charge ratio of a large, complex sample of peptides was measured, the resulting signal would be highly convoluted as many peptides have the same or a very similar m/z. In order to minimize overlapping signals and to get further information on the molecules measured, mass spectrometers are coupled with a previous separation device based on orthogonal (ideally) physical and chemical properties. Typically, the peptides are first separated using liquid chromatography (LC), where molecules are separated and gradually eluted at a certain retention time range, e.g., based on their polarity in the commonly used reversed phase (RP) chromatography. More recently, ion mobility (IMS) has become widely accessible in commercial mass spectrometers as an extra dimension of separation, where ionized molecules are continually separated based on their shape and size before being analyzed in the MS. For instance, in LC-IMS-MS [6][7][8] the retention time and mobility dimensions are recorded in addition to m/z. Figure 1 shows the experimental workflow in the left column.

**Figure 1:**
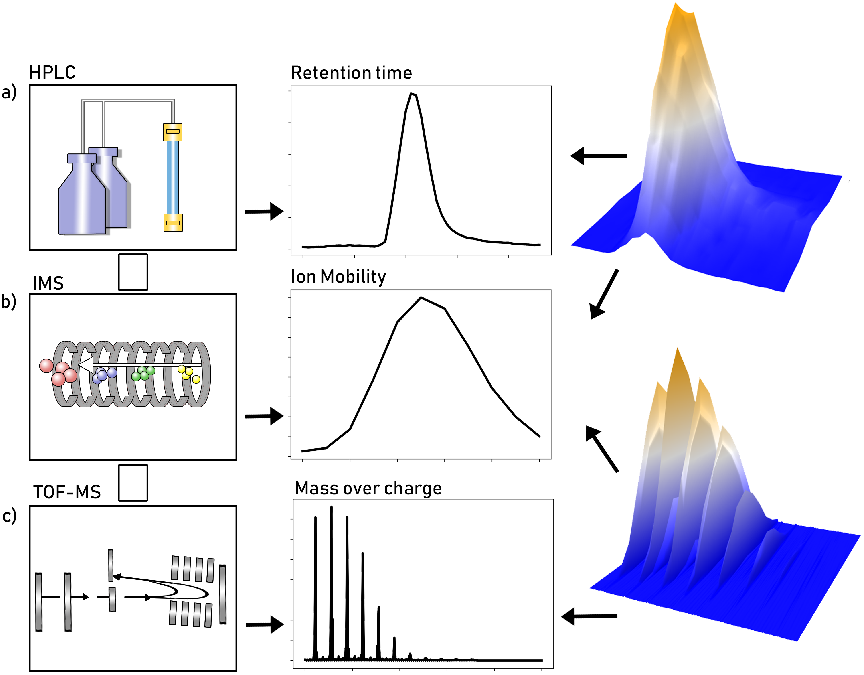
Experimental workflow in an LC-IMS-MS setup and resulting signal of interest. **a)** Chemical separation by high-performance liquid chromatography (HPLC) and resulting intensity distribution along retention time. **b)** Subsequent ion-mobility separation (IMS) and resulting intensity distribution along the mobility dimension. **c)** Time-of-flight mass spectrometry (TOF-MS) and resulting isotopic pattern, i.e., evenly spaced peaks with a certain distribution of the peaks envelope. The column on the right shows the signal as a function of two variables, drift time and retention time on the top and mass-over-charge ratio and drift time on the bottom. Note the repeated occurrence of these isotopic patterns in the 3D plot on the bottom right.

Finally, in many experimental setups two types of mass spectra are recorded: So-called MS1 spectra of all the ions, typically for localization of signals of interest denominated precursors and MS2 spectra, where selected (or unselected) precursors are fragmented and the obtained ion patterns (fragmentation spectra) are typically used for identification.

The first step in the analysis of MS1 spectra is to identify regions of interest. These signals form characteristic patterns in the raw data. More precisely, molecules produce so-called isotopic patterns [9] that are caused by the occurrence of different isotopes in the chemical elements. Thus, one expects a pattern of evenly spaced peaks, where the distance between consecutive peaks varies inversely according to the charge state of the molecule (i.e., a spacing of ≈ 1m/z for *z* = 1, ≈ 0.5m/z for *z* = 2, etc.).

The distribution of peak intensity within an isotopic pattern depends on the chemical elements present in a molecule and their respective distribution of isotopes. Although it is possible to deal with the resulting combinatorial complexity [10], often simpler approaches that assume a typical chemical composition are used to model the distribution of the peak intensity. A common example is the so-called *averagine* model [11].

Due to the coupling with the separation devices, the signals of interest are expected to occur repeatedly over time with the same pattern along the mass axis. Figure 1 shows an example of a resulting signal in the center and right column.

### Problem statement and challenges

The problem to be solved can now be formulated as follows: Given MS1 spectra of a high-throughput setup with additional dimensions of separation, classify whether the signals belong to a region of interest or not.

On top of the signal processing challenge, another technical problems arises: By introducing additional dimensions of separation (such as LC and IMS), the sizes of single data sets increase. Combined with a growth in the number of data sets measured, the storage used for mass spectrometry data has thus grown tremendously over the past years (cf. Figure 2) and is expected to grow further. Consequently, dealing with mass spectrometry raw data will eventually make the usage of Big Data technologies necessary. This means that data will be stored and processed in a distributed manner, which in turn restricts the algorithms applicable.

**Figure 2:**
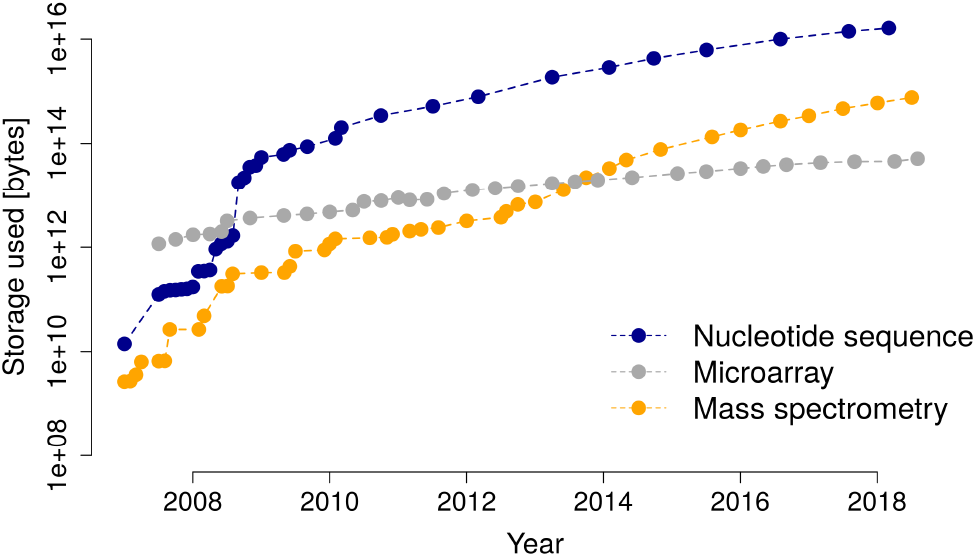
Size of recorded data at the European Bioinformatics Institute (EMBL-EBI) over time for different platforms in life sciences. Mass spectrometry and other data has shown an exponential growth. Plot recreated after [33].

### Locality-sensitive hashing

An often used Big Data method for the comparison of high-dimensional data is locality-sensitive hashing (LSH) [12][13]. In particular, it is a generic algorithm for finding similar pairs of data points (by some measure) in linear runtime. However, this reduction of runtime comes at the prices that the algorithm is probabilistic in nature.

Owing to its wide applicability, the technique is widely used in different fields, including image retrieval [14], pattern recognition [15], and genome analysis [16][17].

### Related work

In the particular context of mass spectrometry, LSH has been used for looking up peptide sequences in databases [18][19], to cluster different spectra for MS1 spectra on LC-MS data [20], and for fast database lookup on MS2 spectra [21].

Previous approaches to signal detection in mass spectrometry raw data either rely on assumptions concerning the isotopic distribution [22] or are based on deep learning [23] and thus lack interpretability.

Concerning the processing of larger data sets, established tools like MaxQuant [24] partially bypass large data sizes by using only parts of the data, i.e., by only looking at every 4th spectrum by default [25].

To the best of our knowledge, signal classification by means of LSH for mass spectrometry raw data has not been treated publicly.

## Results

### Approach

Our approach exploits the fact that all signals from a given region of interest are expected to be similar to each other [25], while noise is assumed to be much more random. Thus, classification of the signal is achieved by deciding whether there are similar signals present. Finding similar objects is achieved by locality-sensitive hashing, as it allows to leverage parallel computation.

The overall scheme of the approach, see Figure 3 for illustration, is the following: A mass spectrometry raw data set is considered to be a set of mass axes for each possible replicated measurement, that is, each retention time for LC-MS data or each retention time and mobility measurement for LC-IMS-MS data, respectively. These mass axes are then cut into small intervals, called windows. For each window several hash values are computed. If two windows have the same hash value they are said to collide. The classification into “true” signal and noise used the following criterion: If a peak lies within a window that collided with any other window it is considered “true” signal, otherwise noise. To facilitate the lookup of collisions, a second map structure is used that maps hash values to their respective number of occurrences.

**Figure 3:**
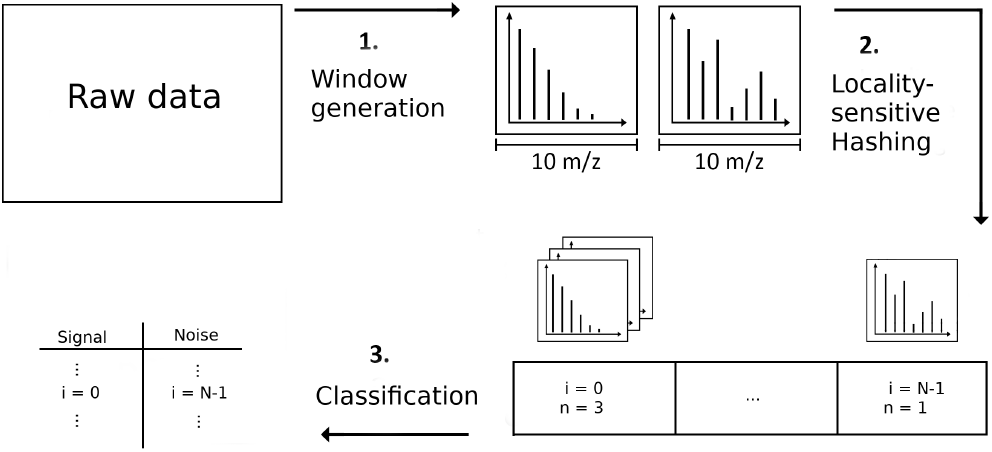
Schematic overview of the approach: Short intervals (windows) of the several mass axes are considered as smallest building blocks of the data set and generated from the raw data. Then for each window the (several) hash functions h map into the hash buckets. Finally, if more than one window is mapped in a given hash box, all windows inside the box are considered “true” signal.

### Advantages of the approach

As the hash function of each window can be computed and checked independently of the other windows, the algorithm is embarrassingly parallel in nature. By using an augmented LSH with *n* AND connectives and *m* OR connectives, the algorithm allows for tuning of false-positive and false-negative rates.

In particular, our approach assumes no model or distributions for the signal shapes, only similarity. Thus, we are able to distinguish different isotopic patterns as true signal, regardless of the composition, size and charge of the ionized molecule.

### Signal classification capability

Figure 4 shows the receiver operating characteristic (ROC) of a classification task on synthetic data. By varying the amplification parameters *m* and *n*^1^ of the LSH, different types of discrimination can be achieved, depending on the goal of the user^2^.

**Figure 4:**
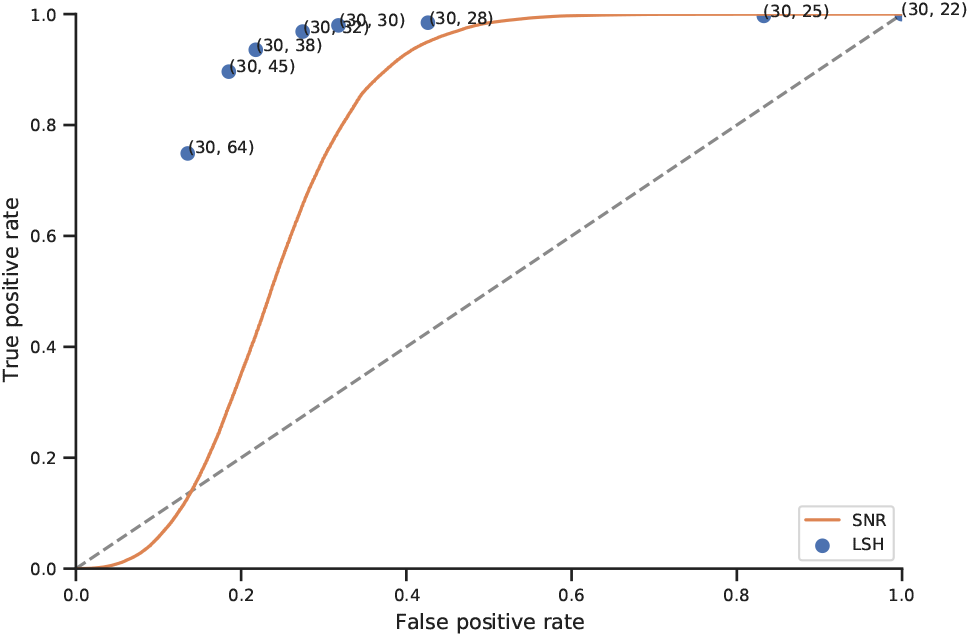
Receiver operating characteristic. Single points mark the results of our approach and the tuples denote the number of AND and OR amplifications used. The solid line shows the performance of the intensity-threshold approach and the dashed line the results of random guessing.

The achieved performance is in accordance with expected behaviour when comparing the implied similarity thresholds in Figure 5. When setting a low threshold, e.g., (*m, n*) = (30,22) (corresponding to the blue line in Figure 5), almost all windows are considered signal, resulting in a true-positive and false-positive rate of almost one.

**Figure 5:**
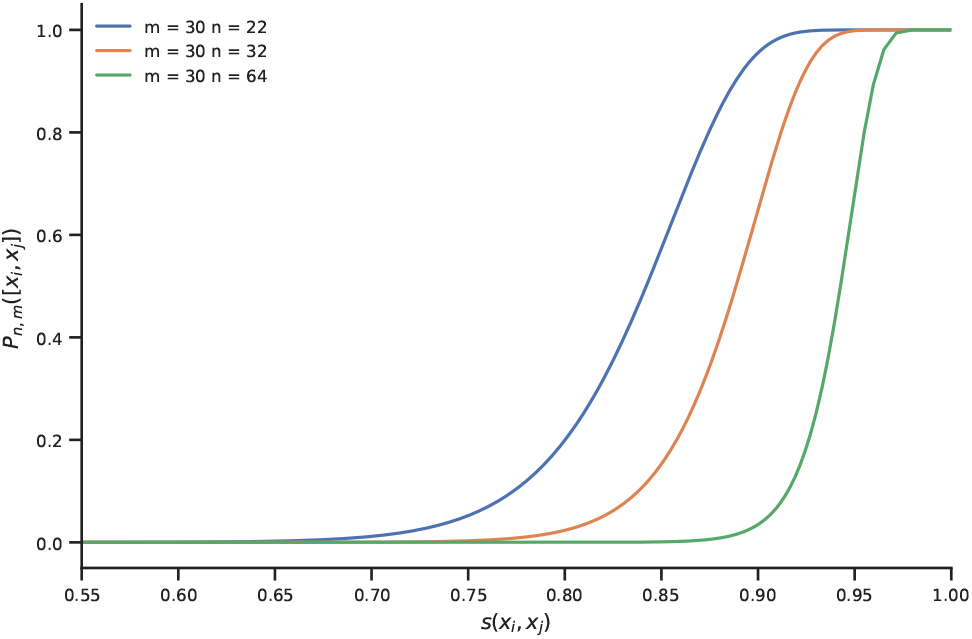
Controlling typical similarity of pairs: Probability *P_m,n_*([*x_i_, x_j_*]) to retrieve a pair for several combined hashes as a function of similarity *s*(*x_i_, x_j_*). By appropriate choice of *m* and *n*, found pairs have a high probability of having at least a certain similarity. Note that the x-axis starts at *s*(*x_i_, x_j_*) = 0.55, for values smaller than that all curves are almost zero.

A stricter discrimination with (*m, n*) = (30, 32) improves the performance significantly. This in turn indicates that the typical similarity measure of the data is in the area where the orange line in Figure 5 has a high slope.

Using a very high threshold, e.g., (*m, n*) = (30, 64) (corresponding to the green line in Figure 5), some windows are lost, resulting in a lower true-positive rate. As the false-positive rate shrinks as well, this setting may be employed as a strong filter to reduce large data sets.

The performance of our approach was compared to the common approach of filtering signal by an intensity threshold. While the overall behavior of its ROC curve is similar to the one of our approach, the performance is considerably better.

Several synthetic data sets with different noise characteristics were created and all main findings persisted, see Supplementary Information.

### Scalability

Figure 6 shows the scalability of the approach for different choices of m and n on parts of a measured data set. In this double-logarithmic plot, the wall-clock time of the implementation roughly follows a linearly decreasing function of the number of threads used. This shows that the implementation scales out well.

**Figure 6:**
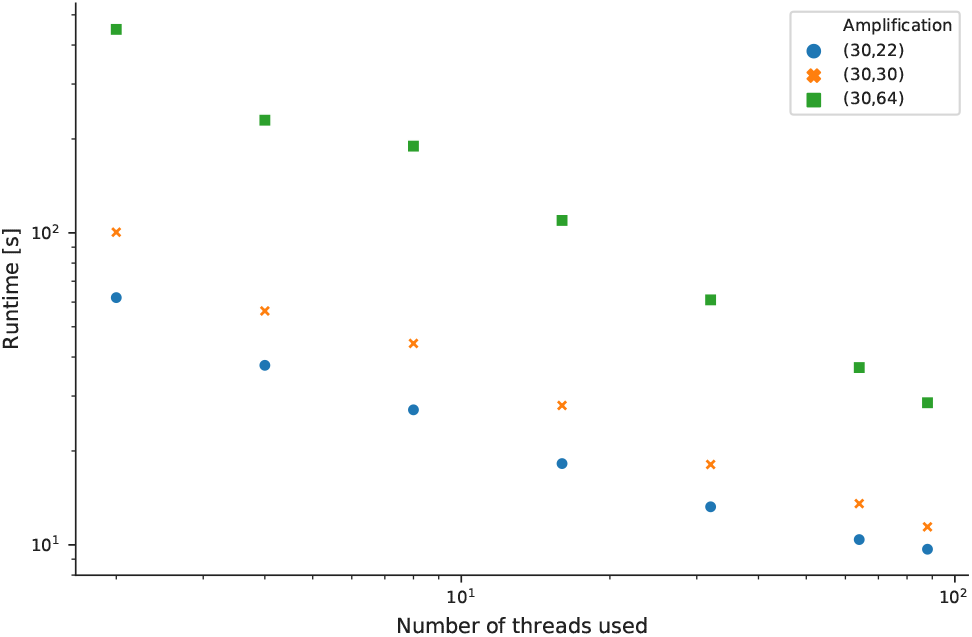
Scalability for different *m* and *n*: Wall-clock time of the implementation in seconds as a function of the number of threads used. Note the logarithmic scales on both axes. The approximately linear trend shows that the implementation scales out well. See text for the setup used.

## Discussion

While the evaluation on synthetic data showed that locality-sensitive hashing could be used in a promising way to detect signals in mass spectrometry raw data, two challenges are expected when applying the method to real world data.

First, in some mass spectrometry setups so-called chemical noise is present. These are signals that originate in chemical impurities in the measurement workflow and are characterized by repeated occurrences. Thus, the chemical noise is self-similar and would be classified as a “true” signal accordingly by this approach.

Finally, signals with a low relative intensity to noise peaks could be lost, as the similarity measure on which the LSH is based drops.

The inability to remove noise peaks from within an isotopic pattern is typically mitigated by the fact that subsequent processing steps like feature finding can take the multidimensional signal shape into account which facilitates the removal of remaining noise peaks.

Testing the approach on real world data is desirable, but we could not find a gold-standard ground-truth peak-level annotation of data.

## Conclusions

Due to the rapidly growing amount of mass spectrometry data, the analysis of mass spectrom-etry raw data could greatly benefit from Big Data methods, most notably implying distributed data storage and highly scalable algorithms.

In this study we showed that locality-sensitive hashing is a desirable approach for signal classification in mass spectrometry raw data. It allows for scalability and provides an approach to signal classification that has a strong focus on self-similarity rather than model assumptions as an intrinsic property of the data.

We propose an implementation using a Big Data framework, such as Apache Spark [26], to facilitate testing on many large data sets from different types of mass spectrometry measurements.

## Methods

### Mass axis windows

For the types of mass spectrometry data considered here, the setup of the mass spectrometer gives rise to a hierarchical structure of the data. In the case of LC-IMS-MS, data is continuously acquired as individual scans across the three dimensions. For each retention time several mobility bins are recorded and in turn for each mobility bin the full spectrum along the mass axis is acquired. In order to allow detection of smaller regions of interest, the algorithm works on short compact subsets (intervals) of the mass axis, henceforth referred to as *windows*. One such window, represented by a list of tuples (m/z,*i*), is the single datum considered by the algorithm for similarity search.

### Window generation

The set of windows is created by dividing all recorded mass axes into windows. In order to avoid missing an isotopic pattern by distributing it into two windows, a second set of windows that is offset by half a window length is created, such that the mass axes are covered in an overlapping fashion.

The length of the windows should be wide enough to capture a whole isotopic pattern, if the windows are applied in an overlapping fashion and small enough such that the chance of having several isotopic patterns in a window is small. For an assumed pattern length of up to 5Da a window length of 10Da is considered useful.

### Binning

The calculation of a window’s hash value requires the mass axis in a binned form with equally spaced bins. Finding an optimal binning scheme is by no means trivial: A very fine binning resolution makes the algorithm less robust and increases the computational cost, while a very coarse binning resolution yields an increased loss of information. A binning resolution of 0.1m/z was considered suitable as, on the one hand, it still resolves isotopic pattern with up to charge state five (corresponding to a spacing of 0.2m/z) and, on the other hand, keeps the computational load feasible.

### Locality-sensitive hashing

#### Used similarity measure

Two windows *W_i_* and *W_j_* shall be considered similar when their mass spectra have the same shape but not necessarily the same overall scale. Therefore, the similarity function *s*(*W_i_, W_j_*) used is the cosine similarity

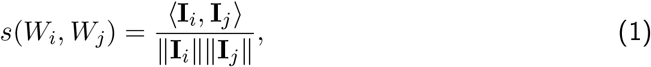

where **I** denotes the intensity array of a binned window, 〈·, ·〉 the standard scalar product and ║·║ the Euclidean norm. As a direct consequence from the linearity of the scalar product and norm, the cosine similarity is scale invariant:

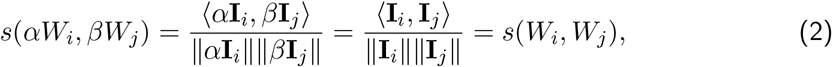

for *α, β* > 0. Thus, the findings of the algorithm are independent of the absolute intensity values.

#### Hash function and amplification

The appropriate family of hash functions for the cosine similarity is the random projection hashing [27]. The hash function is given by *h*(·) = sign(〈·,*r*〉) with the components of the vector *r* being random samples from a standard normal distribution.

To control for false positives or false negatives, an augmented LSH with *n* AND connectives and *m* OR connectives is used. This means that instead of computing a single hash function *m* × *n* hash functions are calculated and divided into *m* arrays of length *n*. Two objects *x_i_* and *x_j_* are now considered collided if all *n* hash functions of a block yield the same value. By choosing m and n appropriately, a sharp sigmoid shape of the collision probability *P_m,n_*([*x_i_,x_j_*]) can be achieved, which effectively translates into a similarity threshold. Figure 5 shows *P_m,n_*([*x_i_,x_j_*]) for different values of *m* and *n*.

### Intensity thresholding

A very simple and general approach to signal classification is by means of thresholding with respect to a signal-to-noise ratio [28]. If one assumes a global noise estimate, this reduces to a thresholding with respect to the intensity of each peak.

### Data generation

The synthetic data sets consist of windows of length of 10m/z, with a binning resolution of 0.1m/z.

In each set, two types of data were created: Firstly, windows containing no isotopic patterns and noise only. Secondly, windows containing both isotopic patterns and noise. The label of each peak (“signal”, if it is part of an isotopic pattern, “noise_2”, if it is a noise peak in a window that contains an isotopic pattern, and “noise_1”, if it is a noise peak inside a window without an isotopic pattern being present) was stored for later evaluation. In total 18445 windows per data set were created, of which 14107 contained noise only and 4338 contained both signal and noise.

In the different data sets the intensities of the “true” signals were scaled differently, such that the maximum signal peak has an intensity of 1000, 500, 250, 125, 64, or 32, respectively.

#### Modeling noise

The noise signal is assumed to consist of independently sampled peaks that share the following properties: The number of peaks per window, *k*, is given as

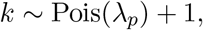

where Pois denotes a Poisson distribution with mean λ_*p*_. Our data was generated using λ_*p*_ = 4.

The location of each peak was sampled uniformly within the window and the intensity value *i* of each peak was sampled from

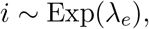

where Exp denotes a exponential distribution with mean 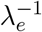. Our data was generated using *λ_e_* = 15.

#### Modeling isotopic patterns

For the “true” signal the averagine model [11] was used. The monoisotopic peak was placed in the middle of the window and the contribution of the next five peaks was considered. In order to model the repeated occurrences of patterns, two copies with all peaks scaled to half intensities were added. Finally, a noise signal, individually sampled according to description above, was added to every true pattern window.

The monoisotopic masses were ranged from 150u to 5000u in steps of 10u. Charge states from 1*e* up to 5*e* were included if the resulting m/z was in the interval [150m/z, 2000m/z].

### Classification by collision

In order to classify windows into “noise” or “signal”, for each window in the data set (several) hash values are computed. Then the number of occurrences of each hash value is counted and all hash values that occurred more then one time are stored in a table, the so-called collision table. A window in question is called “signal” if at least one of its hash values can be found in the collision table, otherwise it is called “noise”.

A single peak in the data set is classified by whether there was a collided window that contains the peak. This is especially important as peaks can be part of several windows due to the overlapping window approach.

### Evaluation

Although the synthetic data provides ground truth labels, a meaningful computation of true-positive and false-negative rates is not straightforward.

Since in our approach all peaks in a window will be assigned the same label, the following problem is caused: When signal and noise is present in a window, peaks with label “noise_2” will be considered “true” as well, which would result in a high false classification rate. For work on real world data however, the inability to remove noise peaks among isotope patterns is typically cured by the fact that subsequent processing steps like feature finding can take the multidimensional signal shape into account, which facilitates the removal of remaining noise peaks.

Thus, in order to learn about the false classification rate of actual interest, peaks with label “noise_2” were not considered for the computation of classification rates both for the intensity thresholding and our approach.

The presence of peaks with label “noise_2” is nevertheless important to test whether whole patterns are missed due to noise in the same window.

### Scalability study

For parallel computing, the algorithm was implemented in C++ with multithreading enabled by usage of openMP [29]. The scalability study was performed on a single Ubuntu 20.04 LTS machine with two Intel Xeon Gold 6238 CPUs, each featuring 22 physical cores @2.1 GHz. Overall, enabled hyper-threading allows for the parallel execution of 88 threads. The machine is further equipped with 192 GiB (12x 16 GiB) of DIMM DDR4 @2933 MHz main memory. Our C++ package was compiled using GCC 9.3 and optimization level −O3.

As test data a single MS1 frame of a nanoLC-TIMS-MS/MS (DDA-PASEF [8]) analysis of HeLa whole proteome digest was used and raw data access was enabled by OpenTIMS [30]. HeLa cells were lysed in a urea-based lysis buffer (7 M urea, 2 M thiourea, 5 mM dithiothreitol (DTT), 2% (w/v) CHAPS) assisted by sonication for 15 min at 4°C in high potency using a Bioruptor instrument (Diagenode). Proteins were digested with Trypsin using a filter-aided sample preparation (FASP) [31] as previously detailed [32]. 200 ng of peptide digest were analyzed using a nanoElute UPLC coupled to a TimsTOF PRO MS (Bruker). Peptides injected directly in an Aurora 25 cm x 75 μm ID, 1.6 μm C18 column (Ionopticks) and separated using a 120 min. gradient method at 400 nL/min. Phase A consisted on water with 0.1% formic acid and phase B on acetonitrile with 0.1% formic acid. Sample was injected at 2% B, lineally increasing to 20% B at 90 min., 35% B at 105 min., 95% at 115 min. and hold at 95% until 120 min. before re-equilibrating the column at 2%B. The MS was operated in DDA-PASEF mode [8], scanning from 100 to 1700 m/z at the MS dimension and 0.60 to 1.60 1/k0 at the IMS dimension with a 100 ms TIMS ramp. Each 1.17 sec MS cycle comprised one MS1 and 10 MS2 PASEF ramps (frames). The source was operated at 1600 V, with dry gas at 3 L/min and 200°C, without nanoBooster gas. The instrument was operated using Compass Hystar version 5.1 and timsControl version 1.1.15 (Bruker). All reagents and solvents used were MS-grade.

## Acknowledgements

The authors would like to thank Mateusz Łącki for fruitful discussions and help in development of ideas and Patrick Raaf for his input on Big Data technology.

## Availability of data and materials

Data and code is available at https://github.com/hildebrandtlab/mzBucket

## Ethics approval and consent to participate

Not applicable.

## Consent for publication

Not applicable.

## Authors’ contributions

BS and AH conceptualized the work, KB and DT designed the work, DGZ acquired data, KB and DT analyzed data, KB, DT and TK created new software used in this work, ST, BS and AH revised the work. All authors read and approved the final manuscript.

## Footnotes

1 A plot with more pairs of *m* and *n* can be found in the Supplementary Information. The best parameters still lie on the implied ROC curve.

2 Note that the number of windows in the data set influences the collision probabilities as well.

