## Supplementary Information for "Locality-sensitive hashing enables signal classification in high-throughput mass spectrometry raw data at scale"

### Different parameters for $m$ and $n$

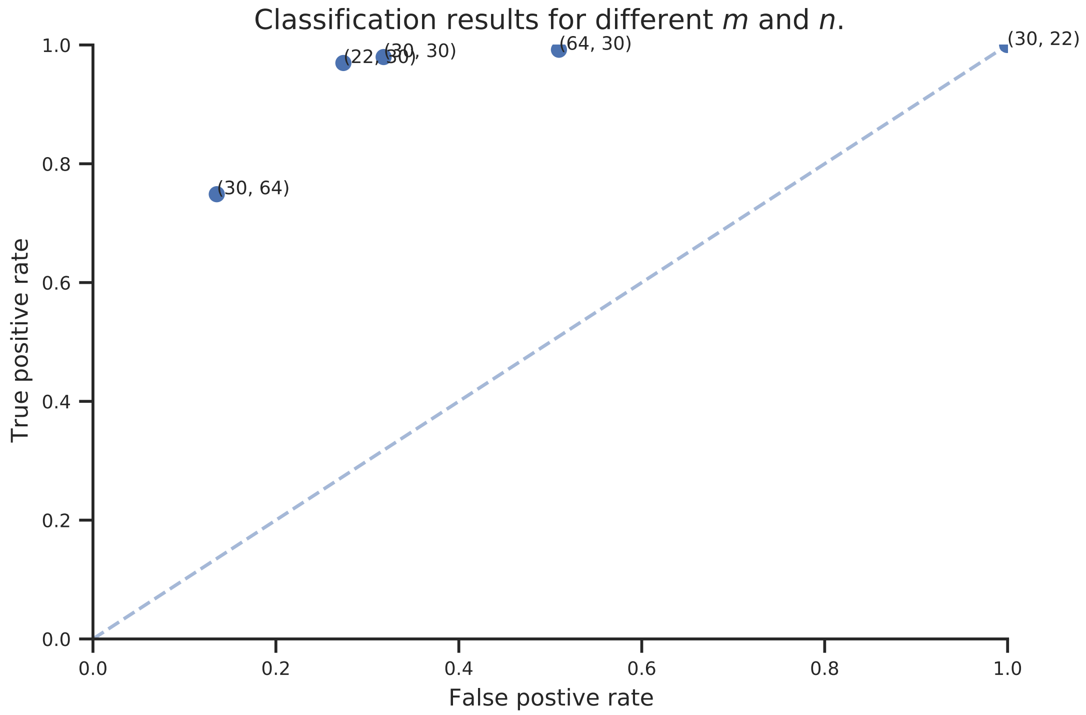


Figure 1: Different parameters for $m$ and $n$. The overall shape of the ROC curve stays the same.

### Different maximum signal intensities in synthetic data

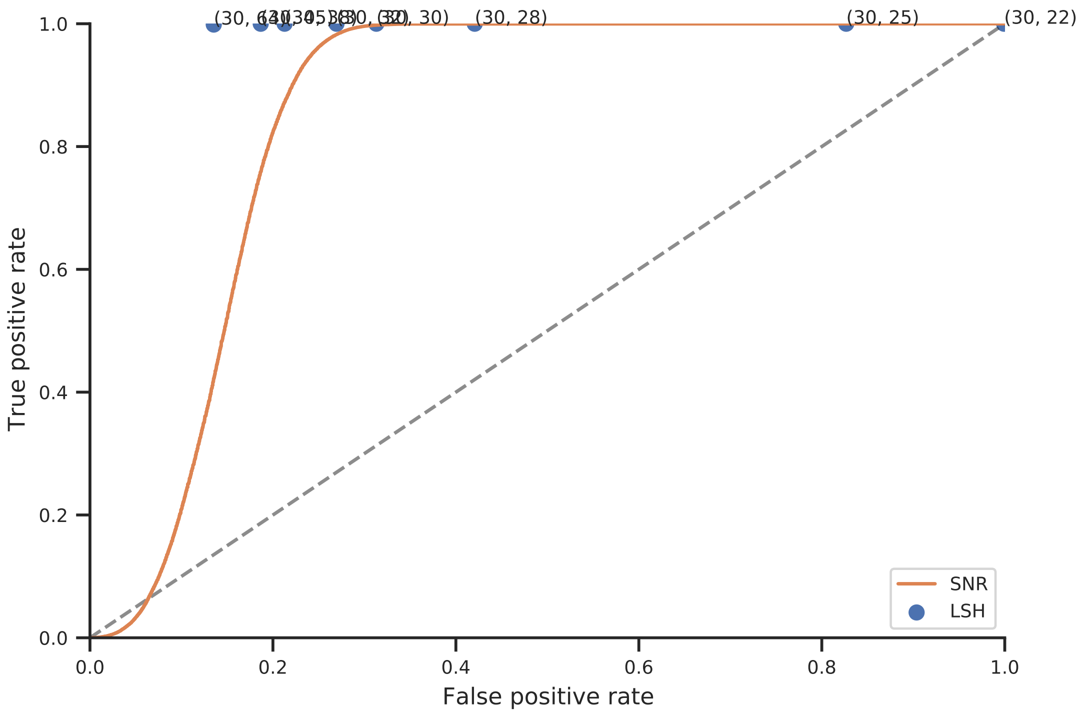


Figure 2: Maximum signal intensity 1000.

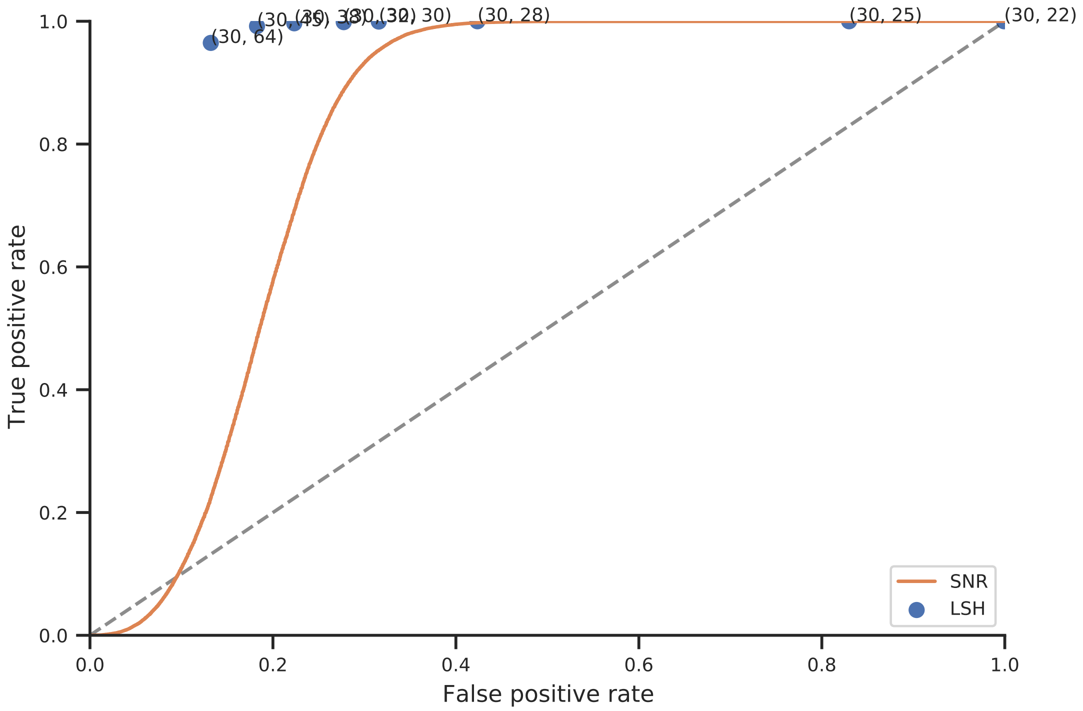


Figure 3: Maximum signal intensity 500.

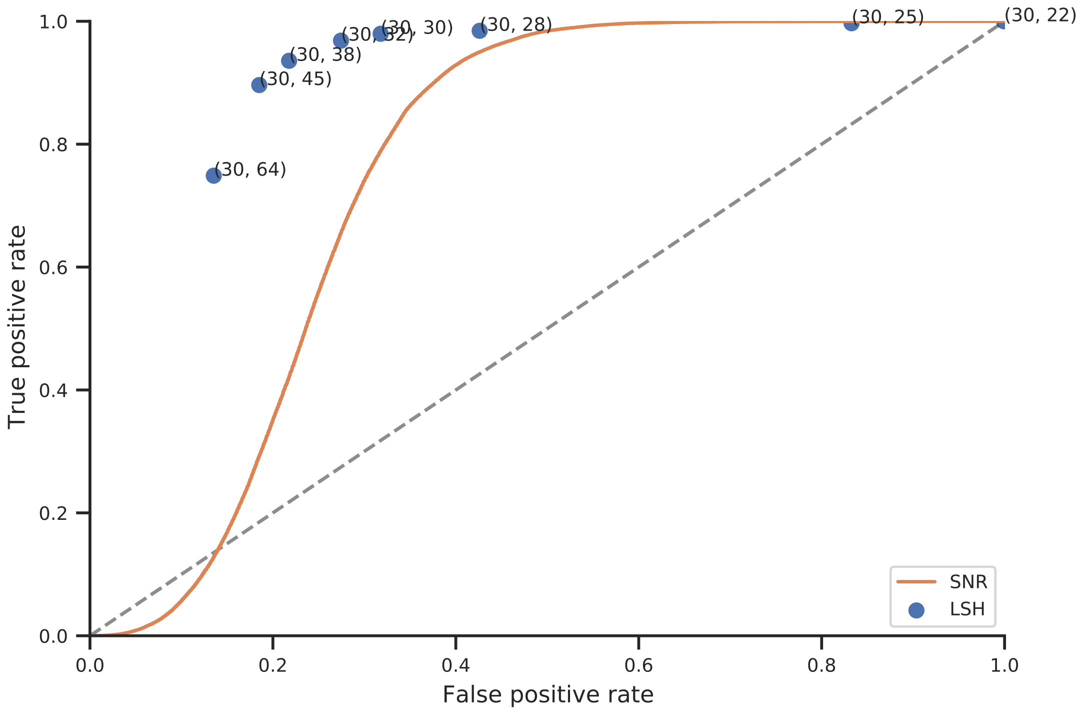


Figure 4: Maximum signal intensity 250.

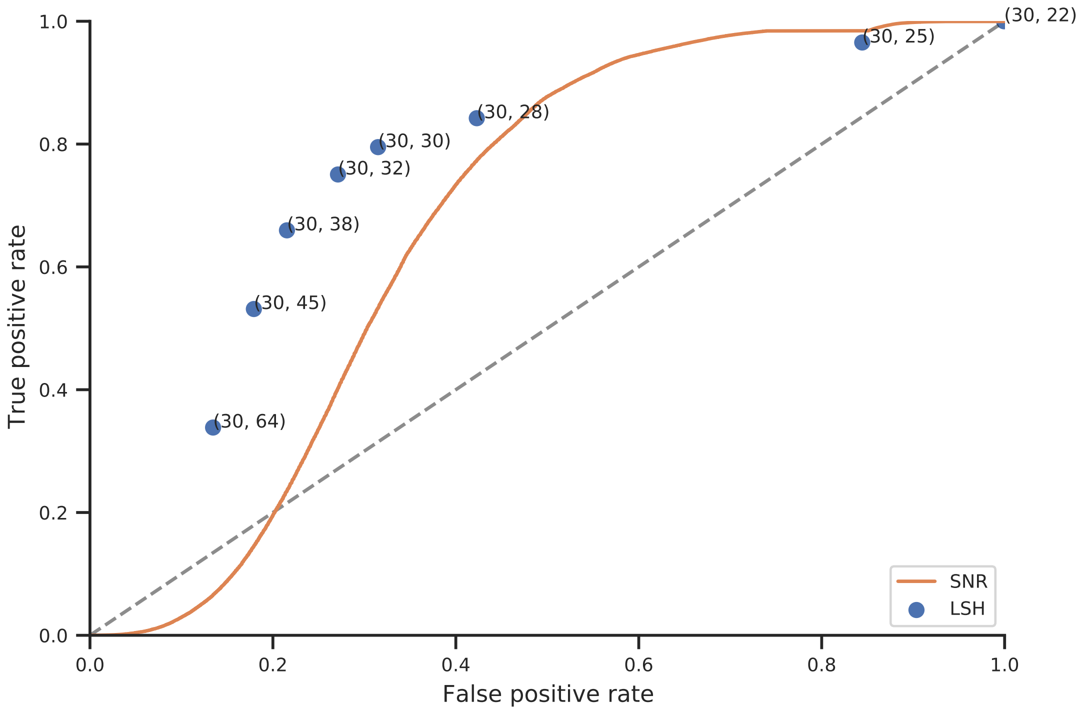


Figure 5: Maximum signal intensity 125.

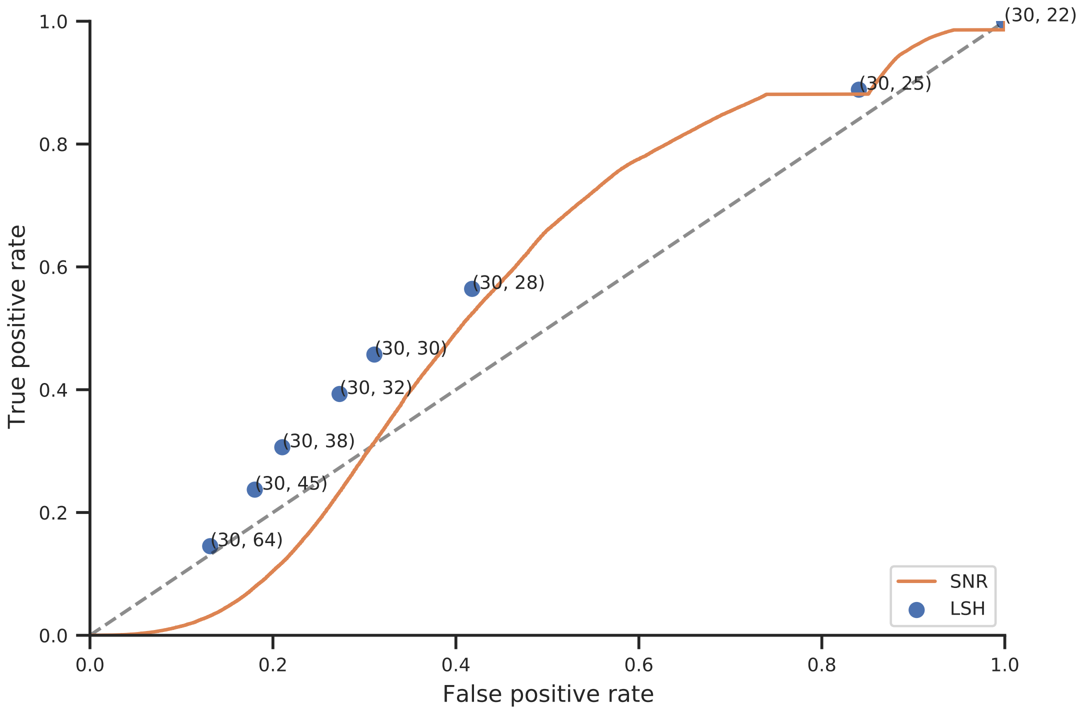


Figure 6: Maximum signal intensity 64.

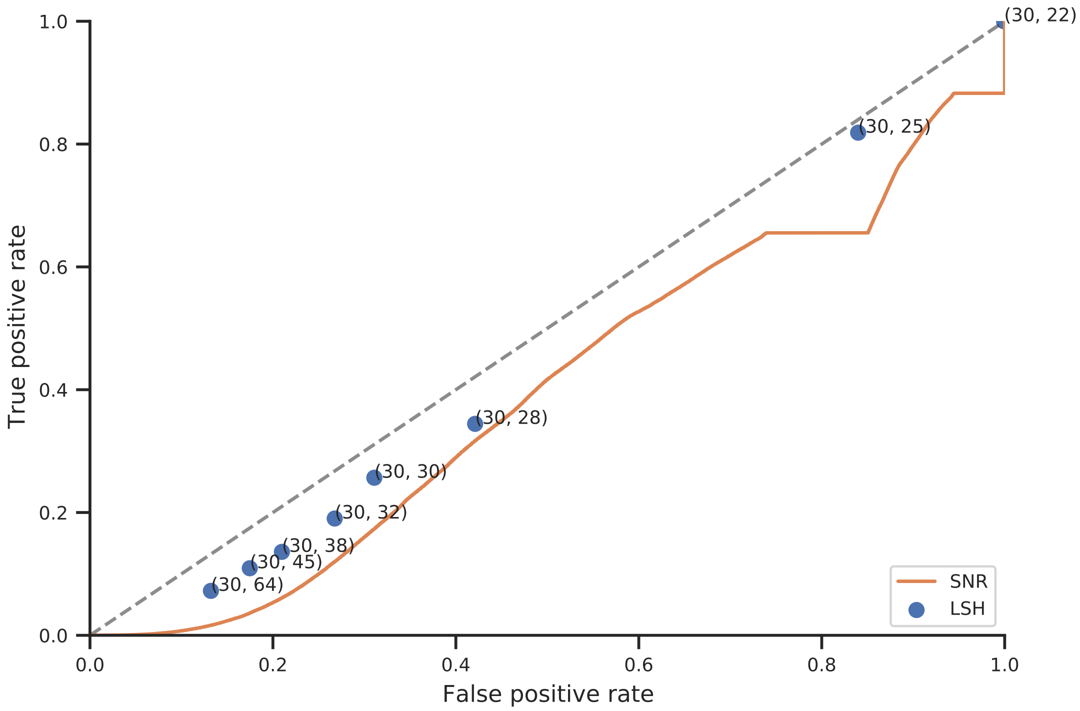


Figure 7: Maximum signal intensity 32.
